# The healthy ageing gene expression signature for Alzheimer’s disease diagnosis: a random sampling perspective

**DOI:** 10.1101/047050

**Authors:** Laurent Jacob, Terence P. Speed

## Abstract

In a recent publication, Sood et al. [2015] presented a set of 150 probe-sets which could be used in a diagnosis of Alzheimer disease (AD) based on gene expression. We reproduce some of their experiments, and show that the performance of their particular set of 150 probe-sets does not stand out compared to that of randomly sampled sets of 150 probe-sets from the same array.

## 2 Correspondence

The first part of the experiments in Sood et al. [2015] builds their signature and assesses its ability to predict chronological age in different settings. This first part involves eight gene expression studies: six from muscle, one from brain and one from skin samples. The skin study was done on Illumina Human HT-12 V3 arrays and all the others on Affymetrix HGU133plus2 arrays. The first muscle dataset involves muscle samples from 15 young and 15 old healthy individuals and is only used to build the signature. The selection process retains probe-sets which are both differentially expressed between young and old samples as measured by limma [Ritchie et al., 2015] and predictive of chronological age in the context of a 5 nearest neighbor classifier, along with other selected probe-sets. The 150 probe-sets selected constitute the healthy ageing gene signature (HAGS) and they are then used in a 5 nearest neighbor classifier to predict the chronological age of samples in the other studies; the study used to select the signature is not used anymore in the rest of their experiments. Sood et al. [2015] use two different protocols to evaluate the prediction performance. For all except the skin data, they use external validation: the samples from one of the muscle studies (Campbell) are used as neighbors to predict the age of the tested samples in the four remaining muscle and the brain study. For the skin study, they use leave one out cross validation (LOOCV). They also use LOOCV on two of the muscle studies and the brain study to produce ROC curves on their Figure 2. They obtain reasonably high AUCs and conclude that their 150 probe-sets are predictive of chronological age regardless of the tissue and platform.

**Figure 2:**
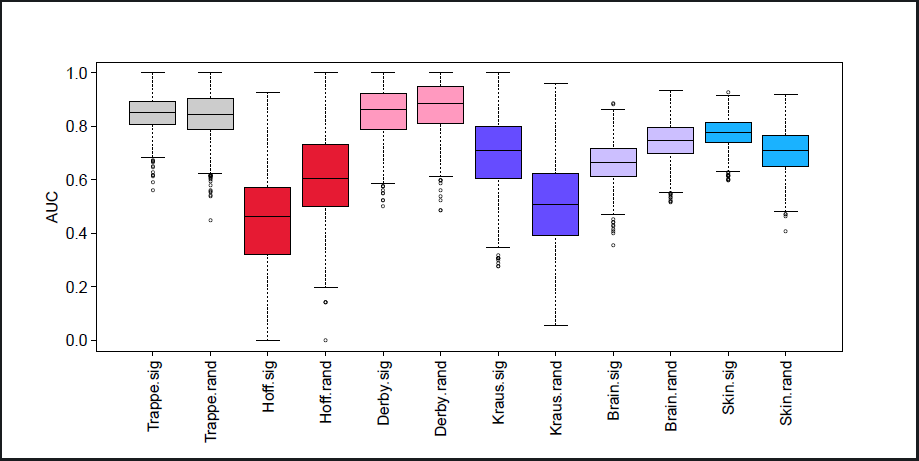
Area Under the ROC Curves obtained by LOOCV of a 5 nearest neighbor classifier over 1000 random selections of 50% of the arrays, using the HAGS probe-sets (.sig suffix) and a new random selection of 150 probe-sets each time (.rand suffix), over the six ageing gene expression studies used in Sood et al. [2015].

In the last part of their experiments, the 150 HAGS probe-sets are used to predict AD status from blood gene expression samples in two cohorts. Gene expression is measured using Illumina Human HT-12 V3 and V4 arrays respectively. Samples are selected within each cohort to make AD status independent of potential confounders such as age, gender or ethnicity. The 150 probe-sets are mapped to Illumina probe-sets. They lead to LOOCV Areas Under the ROC Curve (AUCs) of 0.73 and 0.66.

We reproduce some of the experiments from Sood et al. [2015], showing that the HAGS is indeed able to discriminate between old and young samples in several gene expression studies, and AD from control patients using blood gene expression in two cohorts. We also show that its performance does not stand out compared to that of randomly selected sets of 150 probe-sets from the same array, most of which lead to reasonable performance on these datasets. Finally, our results on random sampling of both arrays and probesets suggest that prediction of either chronological age or AD status on new samples following the same conditional distribution as the ones in these studies would be done, on average, about as well by a random set of genes as by the HAGS. The code used to produce all figures in this report is freely available.

## 3 Comparison of the healthy ageing gene expression signature with random gene sets

We first reproduce the experiments done by Sood et al. [2015] on age data. We obtained the data from the public repositories indicated in the original article. The Affymetrix studies are normalized using RMA as implemented in the Bioconductor affy package. We do not try to reproduce the gene selection process, but use the list provided in the first tab of Additional file 1 of Sood et al. [2015] instead. We extract the list of arrays used in the experiments and the age of the corresponding patients from the third tab of the same additional file. For each of the four muscle studies and the brain study, we measure the AUC obtained by both LOOCV and external validation using the samples from the Campbell dataset as neighbors. For the skin data, we only measure the AUC by LOOCV, as in Sood et al. [2015]. We map the 150 Affymetrix probe-sets to all Illumina probes which match the same gene symbols using the annotate Bioconductor package. Selecting a single Illumina probe for each Affymetrix probe-sets (the one with highest intensity) does not affect the result much so we keep the map using all matching probes, as it does not rely on expression data. The AUCs obtained are represented as green dots on Figure 1, and are generally consistent with the ones obtained by Sood et al. [2015]. The observed differences may be caused by changes in the pre-processing: for example, Sood et al. [2015] use frozenRMA in some cases but we could not understand precisely how. We keep to regular RMA, as it does not seem to affect the performances qualitatively.

**Figure 1:**
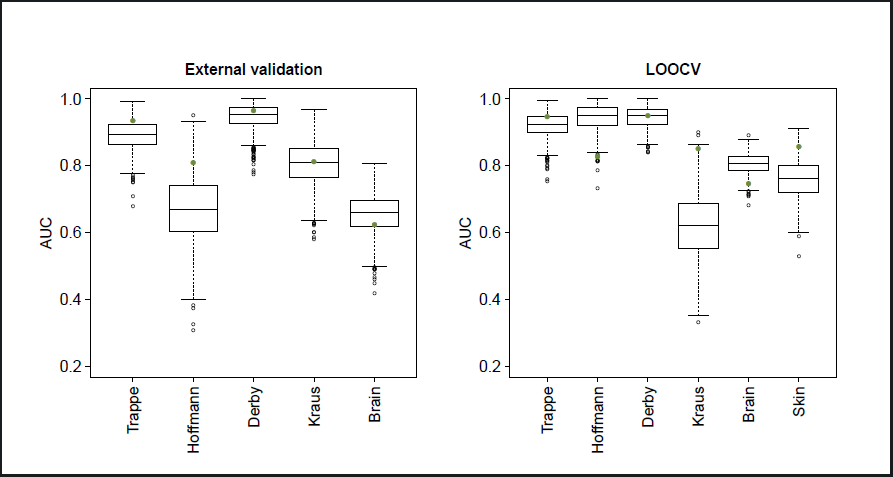
Area Under the ROC Curves obtained by external validation (left panel) and LOOCV (right panel) of a 5 nearest neighbor classifier using the HAGS probe-sets (green dots) and 1000 random selections of 150 probe-sets (boxplots), over the six ageing gene expression studies used in Sood et al. [2015].

We then follow the same protocol using 150 probe-sets randomly sampled from the Affymetrix HGU133plus2 array. For the external validation runs, we sample among the probe-sets that are both present in all studies and in the upper quartile of median absolute deviation in the muscle study that was used by Sood et al. [2015] to select their probe-sets. Sampling among all probes only marginally changes the results. The boxplots on Figure 1 show the distribution of performances obtained on each study across 1000 repetitions of the random sampling. The green dot on each boxplot represents the performance of the 5 nearest neighbor predictor with Euclidean distance using the HAGS, thus quantifying for each study how exceptional its performance is in a context of randomly sampled probe-sets. For most studies the performance of the HAGS does not stand out, and when it does, it is sometimes better and sometimes worse than the median performance of randomly sampled probe-sets.

This first experiment only tells us something about the exceptionality of the HAGS for these six studies. Looking across the studies suggests that the HAGS is not expected to behave differently from 150 randomly selected probe-sets, on average, on new samples from the same distribution^1^. In order to get more evidence on this point, we reproduce the LOOCV evaluation protocol, with further sampling of the arrays: for each of the 1000 repetitions, we perform LOOCV on a randomly sampled 50% of each dataset, with both the HAGS and a new random selection of 150 probe-sets each time.

The results are shown in Figure 2. The comparison of the two empirical distributions on each study tells us which of the HAGS or the random signatures performs better on arrays sampled from the same distribution. For two of the six studies (Trappe and Derby), the distributions are very close. For two others (Kraus and the skin study), the HAGS generally performs better than random signatures. For the last two studies (Hoffman and brain), the HAGS performs worse than random signatures.

Finally, we reproduce the AD status prediction experiments, using the same two random sampling protocols as for the ageing experiments: sampling gene sets and sampling both gene sets and arrays. We stratify our array sampling by status to make sure that the proportion of AD and control status remains unchanged. Further stratifying by age and gender did not change the result. We use the same subset of samples from each cohorts used in Sood et al. [2015], creating two classes by merging the MCI (mild cognitive impairment) and AD status – we refer to this merged class as AD in the remainder of this discussion. For each of the 1000 repetitions, we sample 150 probe-sets from the Affymetrix HGU133plus2 array and map these probe-sets to the Illumina Human HT-12 V3 and V4 probes. We also map the HAGS to the Illumina probes. As for the ageing experiments, when mapping an Affymetrix signature we keep all Illumina probes associated with a gene symbol which is also associated with a probe-set from the signature. Here again, keeping a single Illumina probe for each Affymetrix probe-set does not affect the results. We also obtain the same results by mapping the HAGS to the Illumina probes and then sampling sets of Illumina probes of the same size as the mapped HAGS. External validation is done by using one cohort as the neighbors to predict the status of arrays in the other cohort.

Figure 3 shows the performance obtained by randomly sampling probes only. The performance of the HAGS for discriminating controls from AD patients is not atypical in either of the two cohorts, for either external validation or LOOCV.

**Figure 3:**
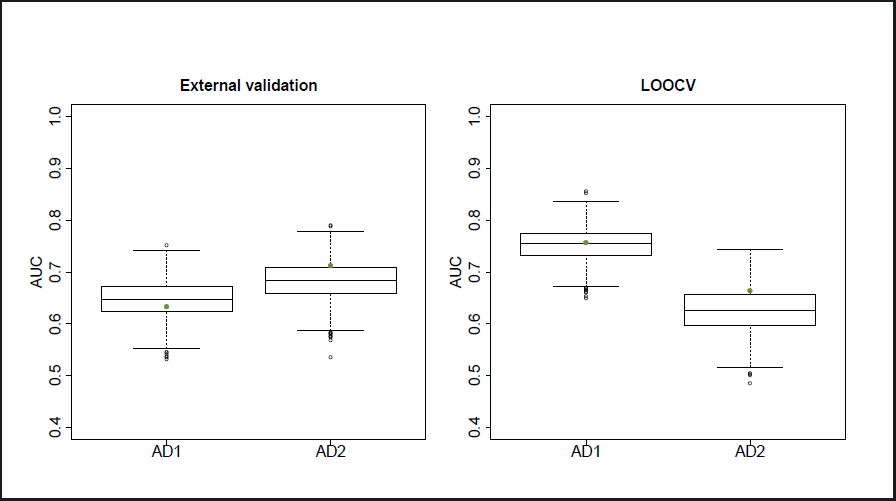
Area Under the ROC Curves obtained by external validation (left panel) and LOOCV (right panel) of a 5 nearest neighbor classifier using the HAGS probe-sets (green dots) and 1000 random selections of 150 probe-sets (boxplots), over the two AD cohorts used in Sood et al. [2015].

Figure 4 further shows that when sampling patients from these two cohorts, the distribution of performances obtained by using the HAGS and by using a different random set of 150 probe-sets for each patient sampling are very similar. This result suggests that when sampling patients from the same distributions as these cohorts, a random set of genes will yield equally good AD status predictions on average as the HAGS.

**Figure 4:**
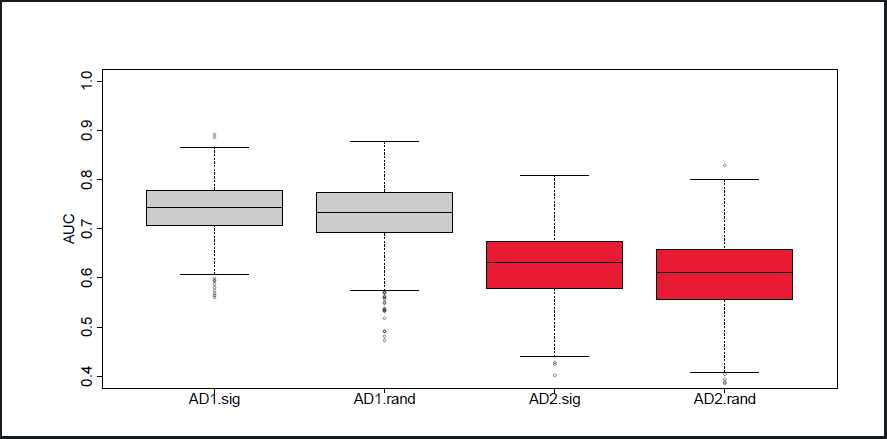
Area Under the ROC Curves obtained by LOOCV of a 5 nearest neighbor classifier over 1000 random selections of 50% of the arrays, using the HAGS probe-sets (.sig suffix) and a new random selection of 150 probe-sets each time (.rand suffix), over the two AD cohorts used in Sood et al. [2015].

The fact that random gene sets perform as well as a set of genes which were selected for their predictive power is not too surprising since the HAGS was selected on a very small number of samples (15 young and 15 old patients) and gene regulation processes make gene expression profiles very highly correlated. It was already noted by Ein-Dor et al. [2005] that sampling from a small set of arrays leads to the selection of different gene expression signatures for breast cancer prognosis. They concluded that “The main lesson is that whenever any arbitrary decision (e.g. choice of training and test set) is taken throughout analysis of the data, one has to generate a large ensemble of the different ways in which this arbitrary decision could be taken, and perform a statistical analysis of the results obtained over this ensemble.” Haury et al. [2011] further studied the stability of feature selection methods, also showing that perturbing the training data leads to the selection of very different signatures, and that small sample sizes are the main reason for this lack of stability. More importantly, they observed in their experiments that a “paired ANOVA test detects no method significantly better than the random selection strategy”. Our finding that randomly selected sets of probes perform as well as the HAGS on average is consistent with their observation. We note two possible reasons for this phenomenon. One is that the phenotypes that are being predicted (25 year old versus 65 year old, diagnosed with AD or MCI versus not diagnosed) may have a strong effect on gene expression, making many sets of genes predictive. The other, regarding AD, is that the HAGS genes were selected as discriminating between young and old healthy patients, making their association with AD status less direct and implying that many other subsets perform equally well at predicting the AD status.

## 4 Implications for using the healthy ageing gene expression signature for AD diagnosis

The results of Section 3 suggest that the HAGS is no better than other sets of 150 probes for predicting the AD status of patients from blood samples, but it does not necessarily imply that the HAGS should not be used for AD diagnosis. Along with most random sets of probes, the HAGS yields reasonably good predictions on both cohorts.

Figure 5 shows the projection of the samples on their first two principal components, with color coded AD status. The first principal component (PC1) explains about 25% of the total variance in both cohorts, and separates the two status rather well. This is consistent with the fact that predicting the AD status from the expression of most 150 probes yields good results, although it may have been the case that only few probes were associated with this PC1. Directly taking the PC1 projection as a predictor of AD status (without using any labeled example) actually yields AUCs of 0.76 and 0.67 on cohorts 1 and 2 respectively. Similar observations can be made when merging the two cohorts and using a linear model to remove the cohort effect (right panels of Figure 5). The AUC obtained is then 0.68.

**Figure 5:**
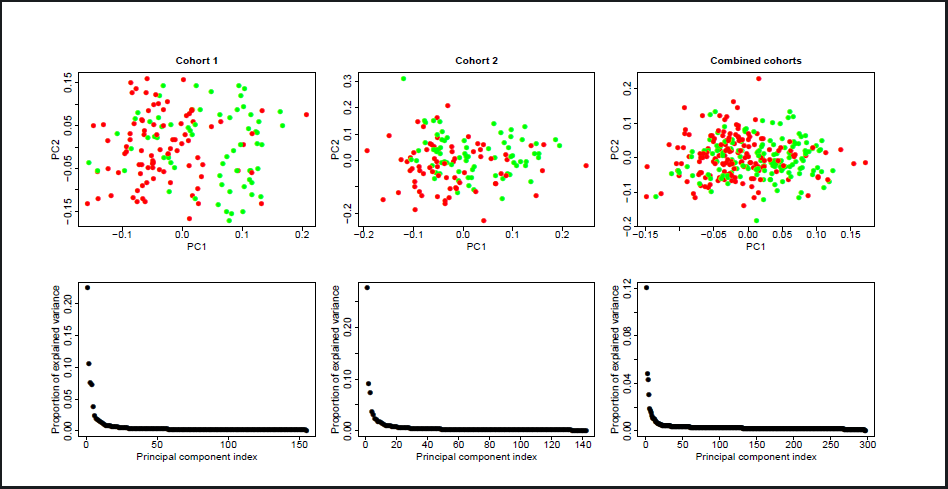
Projection of blood expression samples onto their first two principal components (top) and proportion of variance explained by each principal component (bottom) for cohort 1 (left), 2 (middle) and a combination of both (right).

We also note that 24% of the probes in the first cohort are differentially expressed between AD and control patients (Student’s t-test p-value lower or equal to 0.05)^2^. This proportion is 20% for the second cohort. A uniform sampling of 150 probes would therefore contain 30–40 differentially expressed probes on average.

A possible explanation of the fact that this prediction problem is relatively easy to solve is the presence of an unobserved confounding variable associated with both gene expression measurements and AD status. It is often hard to be entirely sure that no such variable is present, but the authors of Sood et al. [2015] were generally careful to avoid them, in particular blocking by gender, ethnicity and age in their experimental design. Another possiblity is that the problem of discriminating between controls and patients diagnosed with AD from blood gene expression is actually a feasible one because the presence of AD at this stage has a sufficiently strong effect on the overall gene expression. In this case, the question moves to deciding whether a good predictor of current AD status is also a good predictor of future AD status. The latter is arguably a more important objective [Lovestone and Thambisetty, 2009], allowing mass population screenings to detect those at risk, but could prove more difficult than the former as it may be associated with more subtle effects on gene expression. Clinical studies should be able to determine the extent to which the HAGS or other gene expression based predictors are able to diagnose future AD from early blood samples.

## Acknowledgements

The authors thank Anne Biton, Ljubomir Buturovic and Gordon Smyth for their helpful comments.

## Footnotes

1 More precisely, we expect similar performances on average on new samples from the same distribution of the phenotype conditional to the expression of all genes as these six studies.

2 We do not apply a multiple testing correction to these p-values as we are interested in the proportion of discriminating probes for this particular dataset, as opposed to differential expression in a population from which these patients would be sampled.

